# Pharmacological PP2A reactivation overcomes multikinase inhibitor tolerance across brain tumor cell models

**DOI:** 10.1101/2022.05.31.494146

**Authors:** Oxana V. Denisova, Joni Merisaari, Riikka Huhtaniemi, Xi Qiao, Amanpreet Kaur, Laxman Yetukuri, Mikael Jumppanen, Mirva Pääkkönen, Сarina von Schantz-Fant, Michael Ohlmeyer, Krister Wennerberg, Otto Kauko, Raphael Koch, Tero Aittokallio, Mikko Taipale, Jukka Westermarck

**Affiliations:** Turku Bioscience Centre, University of Turku and Åbo Akademi University, Turku, Finland; Institute of Biomedicine, University of Turku, Turku, Finland; Institute for Molecular Medicine Finland, HiLIFE, University of Helsinki, Helsinki, Finland; Icahn School of Medicine at the Mount Sinai, NY, USA; Atux Iskay LLC, Plainsboro, NJ, USA; Biotech Research & Innovation Centre, University of Copenhagen, Copenhagen, Denmark; University Medical Center Goettingen, Goettingen, Germany; Centre for Biostatistics and Epidemiology, University of Oslo, Oslo, Norway; Institute for Cancer Research, Oslo University Hospital, Oslo, Norway; Donnelly Centre, University of Toronto, Toronto, Canada

**Keywords:** DBK-1154, MK-2206, DT-061, NZ-8-061, Dichloroacetate, SMAP

## Abstract

**Background:** Glioblastoma is characterized by hyperactivation of kinase signaling pathways. Regardless, most glioblastoma clinical trials targeting kinase signaling have failed. We hypothesized that overcoming the glioblastoma kinase inhibitor tolerance requires efficient shut-down of phosphorylation-dependent signaling rewiring by simultaneous inhibition of multiple critical kinases combined with reactivation of Protein Phosphatase 2A (PP2A).

**Methods:** Live-cell imaging and colony growth assays were used to determine long-term impact of therapy effects on ten brain tumor cell models. Immunoblotting, MS-phosphoproteomics, and Seahorse metabolic assay were used for analysis of therapy-induced signaling rewiring. BH3 profiling was used to understand the mitochondrial apoptosis mechanisms. Medulloblastoma models were used to expand the importance to other brain cancer. Intracranial xenografts were used to validate the *in vivo* therapeutic impact of the triplet therapy.

**Results:** Collectively all tested ten glioblastoma and medulloblastoma cell models were effectively eradicated by the newly discovered triplet therapy combining inhibition of AKT and PDK1-4 kinases with pharmacological PP2A reactivation. Mechanistically, the brain tumor cell selective lethality of the triplet therapy could be explained by its combinatorial effects on therapy-induced signaling rewiring, OXPHOS, and apoptosis priming. The brain-penetrant triplet combination had a significant *in vivo* efficacy in intracranial glioblastoma and medulloblastoma models.

**Conclusion:** The results confirm highly heterogenous responses of brain cancer cells to mono - and doublet combination therapies targeting phosphorylation-dependent signaling. However, the brain cancer cells cannot escape the triplet therapy targeting of AKT, PDK1-4, and PP2A. The results encourage evaluation of brain tumor PP2A status for design of future kinase inhibitor combination trials.

**Key Points:**

1. Development of triplet kinase-phosphatase targeting therapy strategy for overcoming therapy tolerance across brain tumor models.
2. Identification of interplay between therapy-induced signaling rewiring, OXPHOS, and BH3 protein-mediated apoptosis priming as a cause for kinase inhibitor tolerance in brain cancers.
3. Validation of the results in intracranial in vivo models with orally bioavailable and brain penetrant triplet therapy combination.

**Importance of the Study:** Based on current genetic knowledge, glioblastoma should be particularly suitable target for kinase inhibitor therapies, However, in glioblastoma alone over 180 clinical trials with kinase inhibitors have failed. In this manuscript, we recapitulate this clinical observation by demonstrating broad tolerance of brain cancer cell models to kinase inhibitors even when combined with reactivation of PP2A. However, we discover that the therapy-induced signaling rewiring, and therapy tolerance, can be overcome by triplet targeting of AKT, PDK1-4 and PP2A. We provide strong evidence for the translatability of the findings by orally dosed brain penetrant triplet therapy combination in intracranial brain cancer models. The results encourage biomarker profiling of brain tumors for their PP2A status for clinical trials with combination of AKT and PDK1-4 inhibitors. Further, the results indicate that rapidly developing PP2A reactivation therapies will constitute an attractive future therapy option for brain tumors when combined with multi-kinase inhibition.

## INTRODUCTION

Even though kinase inhibitors have revolutionized cancer therapies, most tumors acquire resistance to kinase inhibitors and their combinations.^1,2^ Especially in cancer types genetically associated with hyperactivation of kinase pathways, such as human glioblastoma, the clinically observed kinase inhibitor resistance is a mechanistic enigma.^3-6^ Acquired therapy resistance develops via two phases - first through adaptive development of a drug-tolerant cellular state, and later, stable resistance that often occurs through acquisition of genetic mutations.^7^ The emerging evidence strongly indicates that the drug-tolerance is initiated rapidly after drug exposure by non-mutational signaling rewiring, often mediated by phosphorylation dependent signaling pathways.^8,9^ Thereby, characterization of the phosphorylation-dependent signaling rewiring events, and kinases/phosphatases controlling the rewiring, can provide novel approaches for targeting the brain tumor relapse at its roots.^10^

Glioblastoma (GB) is the most common primary brain tumor in adults associated with high degree of therapy resistance, tumor recurrence and mortality.^5,11^ Extensive genome-wide profiling studies have established receptor tyrosine kinase RTK/RAS/PI3K/AKT signaling as one of the core altered pathways contributing to GB disease progression.^3,6,12^ AKT pathway fuels aerobic glycolysis,^13^ and GB cells are notorious for employing aerobic glycolysis in energy production and survival.^14,15^ However, targeting of the deregulated AKT and mitochondrial metabolism pathways, even by combination therapies, have achieved dismal clinical response rates in GB.^4,16,17^ In addition to challenges with drug delivery across the brain-blood barrier (BBB) with a number of kinase inhibitors, the failure of kinase targeted therapies in GB is linked to the prevalence of kinase pathway-mediated rewiring mechanisms,^17^ and general apoptosis-resistance of glioblastoma stem-like cells (GSCs).^11^ Also the great intratumoral heterogeneity of GB constitutes a significant therapeutic challenge as the therapies should be effective across cells with different lineage and differentiation status as well as different signaling pathway activities.^11,18^ Tumor suppressor PP2A broadly regulates phosphorylation-dependent signaling and its pharmacological reactivation has in other cancer types shown to impact kinase inhibitor tolerance.^19-21^ However, it is unclear whether pharmacological PP2A reactivation would be able to overcome the kinase inhibitor tolerance across heterogenous human brain tumor models.

## METHODS

### Ethics Statement

In vivo experiments have been authorized by the National Animal Experiment Board of Finland (ESAVI/9241/2018 license), and studies were performed according to the instructions given by the Institutional Animal Care and Use Committees of the University of Turku, Turku, Finland. The animal experiments for this study described in Supplementary Methods.

### Cell culture and reagents

Established human GB cell lines T98G, U87MG, A172, U118, U251, E98-FM-Cherry, patient-derived GSCs, BT3-CD133^+^ and BT12, and human fibroblasts were cultured as described in.^22,23^ Medulloblastoma cell lines DAOY and D283-Med were purchased from ATCC and cultured in Eagle MEM. All cell cultures were maintained in a humified atmosphere of 5% CO_2_ at 37°C. For assays requiring adherent cell, GSCs were cultured on Matrigel (Becton Dickinson) coated plates.

*PPME1* knockout T98G cells were generated as described in.^24^ SV40 small T expressing T98G cells were generated using SV40 small T expressing piggyBac plasmid (pPB-ST, gift from Vera Gorbunova, University of Rochester, NY, USA) by the nucleofection method described in.^25^

UCN-01, AKT1/2 inhibitor, sodium salt of dichloroacetate (DCA) were purchased from Sigma-Aldrich and MK-2206 from MedChemExpress. SMAPs (NZ-8-061, DBK-794, DBK-1154 and DBK-1160) were kindly supplied by Prof. Michael Ohlmeyer (Atux Iskay LLC, Plainsboro, NJ, USA), were dissolved in DMSO and stored at room temperature protected from light.

### RNAi-based knockdown

T98G cells (2 × 10^5^ into 6 well plate) were reverse transfected with siRNAs (Table S3) using Lipofectamine RNAiMAX (Invitrogen) according to the manufacturer’s instructions. On the next day, optimized numbers of cells were re-plated into either 96-or 12-well plates and used for cell viability or colony formation assays, see Supplementary Methods.

### Caspase-3 and -7 activity assay

T98G cells (2.5 × 10^3^) were plated in 96-well plates and allowed to adhere. After 24 hours, cells were treated with the indicated drugs in combination with pan-caspase inhibitor Z-VAD-FMK (10 mM, Promega). After 24 hours, caspase-3 and -7 activities were measured by Caspase-Glo 3/7 assay (Promega) according to the manufacturer’s instructions.

### Long-term growth assay

E98 cells (3 × 10^3^) were plated in 96-well plate. On the next day cells were treated with DMSO, MK-2206 (7 µM), DCA (20 mM), NZ-8-061 (10 µM) alone, or in their doublet or triplet combinations (6-12 wells/condition). Every 3-4 days medium was replaced with fresh media with or without drugs. The confluency of the wells was determined daily using an IncuCyte ZOOM live cell analysis system (Essen Bioscience).

### Immunoblotting

Immunoblotting was performed as previously described.^22^ Primary antibodies: AKT (Cell Signaling, 9272S, 1:1000), phospho Akt S473 (Cell Signaling, 9271, 1:1000), PME-1 (Santa Cruz Biotechnology, sc-20086, 1:1000), phospho PDHE1α S300 (Millipore, ABS194, 1:1000), cleaved PARP1 (Abcam, ab32064, 1:1000), SV40 T Ag Antibody (Pab 108) (Santa Cruz Biotechnology, sc-148, 1:1000), β-actin (Sigma-Aldrich, A1978, 1:10 000) and GAPDH (HyTest, 5G4cc, 1:10 000). Secondary antibodies were purchased from LI-COR Biotechnology.

### Mitochondrial respiration measurement

To assess basal cellular metabolic activity, Agilent Seahorse XF Cell Mito Stress Test (Agilent Seahorse Bioscience) was applied according to the manufacturer’s instructions. Details are described in the Supplementary Methods.

### BH3 profiling

BH3 profiling was performed as previously described.^26,27^ For details, see Supplementary Methods.

### LC-MS/MS analysis of FFPE samples

The LC-ESI-MS/MS analyses were performed on an Orbitrap Fusion Lumos mass spectrometer (Thermo Fisher Scientific) equipped with a nano-electrospray ionization source and FAIMS interface. Compensation voltages of -40 V, -60 V, and -80 V were used. With Orbitrap Fusion Lumos MS data was acquired automatically by using Thermo Xcalibur 4.4 software (Thermo Fisher Scientific). A DDA method consisted of an Orbitrap MS survey scan of mass range 350– 1750 m/z followed by HCD fragmentation for the most intense peptide ions in a top speed mode with cycle time 1 sec for each compensation voltages.

### Statistical analyses

For cell culture experiments, three biological replicates have been performed, and each condition was tested in triplicate, unless otherwise specified. Data are presented as mean ±SD and statistical analyses were carried out using a two-tailed Student’s t-test assuming unequal variances. For *in vivo* experiments, the following statistical tests were chosen depending on the results of the preliminary Shapiro-Wilk test of data normality. Lonk-rank (Mantel-Cox) test was used in survival analysis. These univariate statistical analyses were performed using GraphPad Prism 9 software. P<0.05 was considered statistically significant.

## RESULTS

### Pharmacological reactivation of PP2A synergizes with a multi-kinase inhibitor UCN-01

PP2A is frequently inactivated in GB by non-genetic mechanisms including overexpression of endogenous PP2A inhibitor proteins such as CIP2A, PME-1, SET and ARPP19.^22,28,29^ In a previous study, PP2A reactivation by siRNA-based inactivation of PME-1 was found to sensitize GB cells to several staurosporine multi-kinase inhibitor (MKI) derivatives, including UCN-01.^23^ However the translational impact of these results was questionable as neither the siRNA therapies for brain tumors are not sufficiently advanced, nor does the UCN-01 cross the BBB. To provide potential translational advance, we hypothesized that the recently developed BBB permeable PP2A reactivating compounds (SMAPs)^20,22,30^, could be a pharmacological approach to induce synthetic lethal drug interaction^31^ with UCN-01 in GB cells (Fig. 1A). To test the hypothesis, we directly compared the synergy with UCN-01 and PP2A reactivation by either PME-1 depletion,^32^ or SMAP (NZ-8-061) treatment, on colony growth potential of T98G cells. As shown in Fig. 1B, PME-1 depletion (either by siRNA or by CRISPR/Cas9) or NZ-8-061 did not induce any significant growth defect but induced potent synthetic lethality (SL) with UCN-01. The interaction between NZ-8-061 and UCN-01 was dose dependent and observed by using both compounds at concentrations that showed negligible monotherapy activity (Fig. 1C, D, S1A). Validating the particular potential of PP2A reactivation in kinase inhibitor sensitization,^10^ NZ-8-061 displayed synergistic activity with as low as 0.5-2 µM concentration, that is approximately 10-fold lower concentrations that has been previously shown to be required for monotherapy effects for the compound.^22,30^. NZ-8-061 has been shown in number of publications to directly interact with, and impact PP2A complex composition both *in vitro* and *in cellulo*^30,33,34^. Consistent with these results, and our observations that low micromolar concentrations of NZ-08-61 are sufficient therapeutic effects in drug sensitization, we observed clear evidence for *in cellulo* target engagement of NZ-08-61 with B56 subunits of PP2A by Proteome Integral Solubility Alteration (PISA) assay^35^ from T98G cells treated for with 2 µM of NZ-08-61 for 3 hours (Kauko et al., data not shown). Further, consistently with published data demonstrating rescue of NZ-8-061 effects by overexpression of selective PP2A inhibitor protein SV40 small t-antigen (SV40st)^30,36^, the drug interaction between UCN-01 and NZ-8-061 was abrogated in SV40st expressing T98G cells (Fig. 1E, S1B). Induction of caspase 3/7 activity indicated that the mode of cell death by SMAP+UCN-01 combination was apoptosis (Fig. 1F). To further rule out that the synergy between NZ-8-061 and UCN-01 would be mediated by any potential non-selective targets of NZ-8-061, we used SMAPs DBK-794 and DBK-1154 derived from dibenzoapine tricyclic family, i.e. chemically different from NZ-8-061 (Fig. 1G). Both DBK-794 and DBK-1154 were originally used to demonstrate direct interaction between SMAPs and PP2A, and for mapping of their interaction region.^30^ Importantly, these chemically diverse PP2A reactivators all resulted in identical drug interaction with UCN-01 (Fig. 1H, S1C).

**Figure 1.**
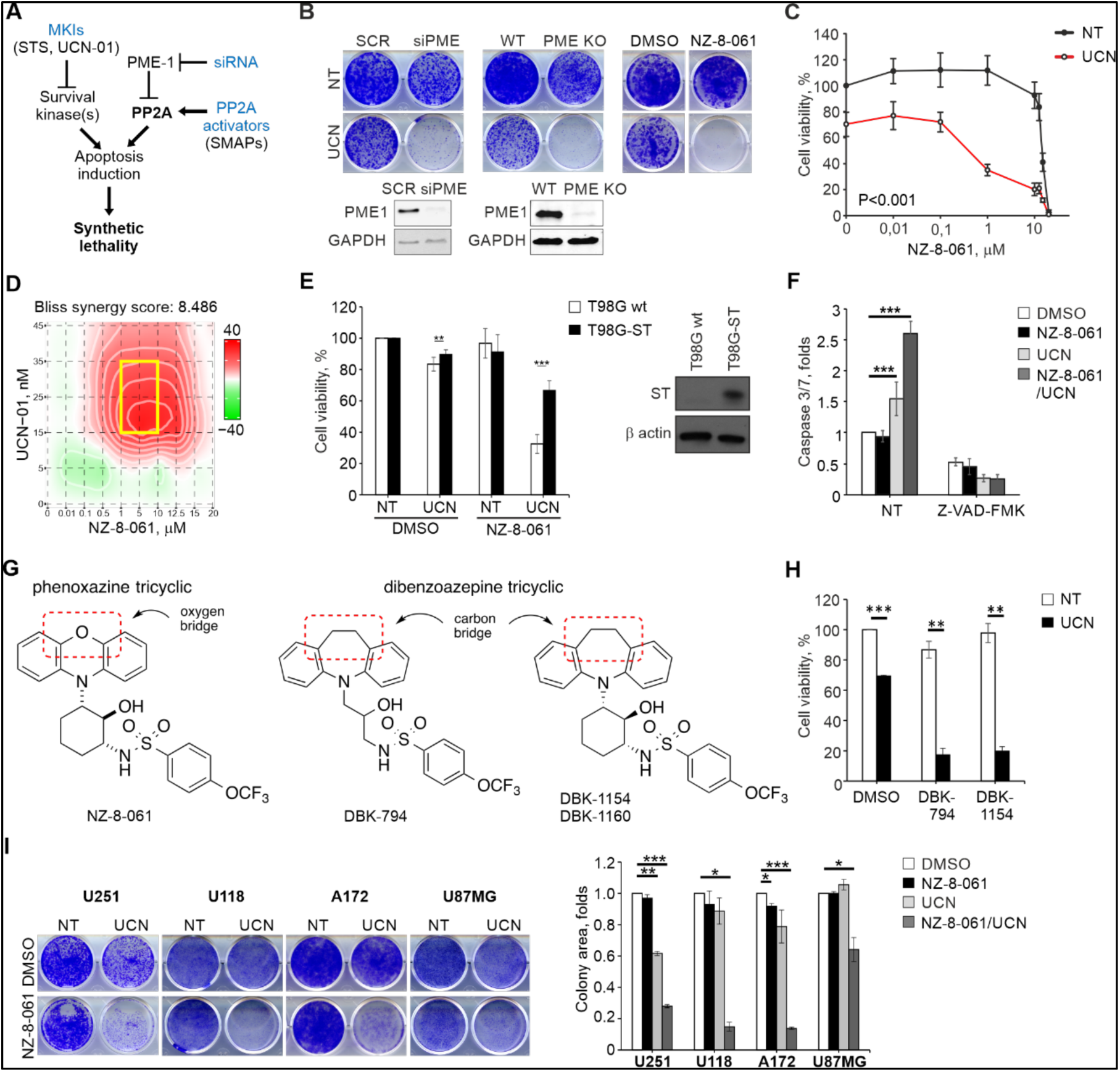
PP2A reactivation and UCN-01 exert a synergistic effect in GB. **A)** Schematic illustrating PP2A reactivation predisposed to MKI-induced SL in GB. **B)** Representative images of colony formation assay in T98G cells under PME-1 deletion (siRNA or CRISPR/Cas9) or NZ-8-061 treatment. Cells were treated with 25 nM UCN-01 (UCN) or left untreated (NT). Immunoblot analysis of PME-1 (lower panel). **C)** Viability of T98G cells treated with increasing concentration of NZ-8-061 either alone or in combination with 25 nM UCN-01 (UCN) for 72 h. ***P<0.001, Student’s *t*-test. **D)** Synergy plot showing the most synergistic area (yellow box) between NZ-8-061 and UCN-01 in T98G cells. The Bliss synergy score is calculated over the whole dose-response matrix. **E)** Viability of T98G wt and SV40 small t-antigen-expressing (T98G-ST) cells treated with 25 nM UCN-01 (UCN) and 8 µM NZ-8-061, alone or in combination for 72 h. Immunoblot analysis of SV40 small t-antigen (right panel). **P<0.01, ***P<0.001 Student’s *t*-test. **F)** Caspase 3/7 activity in T98G cells treated with 8 µM NZ-8-061 alone or in combination with 25 nM UCN-01 (UCN) under caspase inhibitor Z-VAD-FMK (20 µM) for 24 h. ***P<0.001, Student’s *t*-test. **G)** Structures of two different classes of SMAPs. **H)** Viability of T98G cells treated SMAPs, 10 µM DBK-794 and 5 µM DBK-1154, alone or in combination with 25 nM UCN-01 (UCN) for 72 h. **P<0.01, ***P<0.001, Student’s *t*-test. **I)** Representative images (left) and quantified data of colony formation assay (right) in U251, U118, A172 and U87MG cells treated with 8 µM NZ-8-061 alone or in combination with UCN-01 (UCN; 200 nM, 25 nM, 50 nM and 500 nM, respectively). n=2 independent experiments, *P<0.05, **P<0.01, ***P<0.001, Student’s *t*-test.

Together with identical synergy observed by genetic PP2A reactivation (Fig. 1B)^23^, target engagement data, rescue with small-t overexpression (Fig. 1A, S1B), and induction of synergy with non-toxic low micromolar SMAP concentration (Fig. 1D), the use of SMAPs with different chemistry mitigate concerns that the SMAP effects would be related to potential non-selective effects reported using toxic (10-30 µM) concentrations of NZ-8-061 (a.k.a. DT-061).^37^ Additionally, the drug interaction was validated across multiple GB cell lines (Fig. 1I, S1C). Importantly, synergy between UCN-01 and NZ-8-061 was not observed in non-cancerous fibroblasts providing evidence for cancer selectivity of the drug interaction (Fig. S1C, D). The synergistic drug interaction in GB cells was also seen in hypoxic environment, which is a common resistance mechanism in GB (Fig. S1E).

### Triplet kinase/PP2A targeting is required for cytotoxic cell killing across heterogenous brain tumor cell lines

Results above demonstrate strong synergistic activity between BBB-permeable pharmacological PP2A reactivation and multi-kinase inhibition by UCN-01. However, as UCN-01 targets approximately 50 different kinases at nanomolar concentrations,^38,39^ it was necessary to identify kinases that are specifically involved in SL phenotype observed in combination with PP2A reactivation. To facilitate this, we developed a generalizable target kinase screening strategy designated as Actionable Targets of Multi-kinase Inhibitors (AToMI). Detailed description of the AToMI approach can be found from a separate publication.^24^ Based on AToMI screening, both PI3K/AKT/mTOR pathway, as well as mitochondrial pyruvate dehydrogenase kinases (PDK1 and PDK4) were identified as candidate UCN-01 targets that preferentially synergized with either pharmacological or genetic PP2A reactivation.^24^ Notably, immunoblot analysis revealed constitutive, but highly heterogeneous AKT and PDK1-4 activity across most of the brain tumor cell models used in this study (Fig. S2A). The validation results^24^ across three established GB cell lines (T98G, E98, U87MG) and two patient-derived mesenchymal type GSC lines (BT-CD133^+^, BT12) showed that either genetic (PME-1 inhibition), or pharmacological (NZ-8-061 and DBK-1154), PP2A reactivation sensitized the cells to selective AKT (MK-2206 or AKT1/2i) or PDK1-4 inhibitors (DCA or lipoic acid).^15,40^ However, illustrative of the challenge with heterogeneity of GB cell therapy responses, maximal inhibition of cell viability with kinase inhibitor/PP2A reactivator doublet combinations (NZ-8-061+MK-2206, or NZ-8-061+DCA) was highly variable across the cell lines, and in most cases only reached cytostatic effect i.e., about 50% inhibition (Fig. S2B). As cytostasis is generally not considered to be sufficient for durable therapeutic response,^41^ these results indicate that unlike suggested for other cancer types,^19^ PP2A reactivation combined with either AKT or PDK1-4 inhibition cannot be used as a general strategy to kill heterogeneous GB cell populations.

Therefore, we decided to combine both kinase inhibitors together with PP2A reactivation as a triplet therapy (NZ-8-061+MK-2206+DCA). The rationale behind the triplet combination was that non-genetic signaling rewiring induced by single and doublet therapies^8,9^ could be avoided by simultaneous targeting of two major kinase signaling nodes and lowering of the serine/threonine phosphorylation by PP2A activation. To be able to assess GB cell responses to mono, doublet, and triplet therapies both quantitatively and qualitatively, we performed an Incucyte long-term confluency analysis in E98 cells treated with drugs twice for one week, with one week drug holiday in between (Fig. 2A). Consistent with short-term viability assay results,^24^ the E98 cells displayed cytostatic responses to monotherapies during the first 6-day dosing period (Fig. 2A). However, the long-term data confirmed that E98 cells fully escaped all these monotherapy effects. Further, although doublet combinations were found to be more efficient than monotherapies, the cells were able to regain their proliferation after the drug wash out, indicating for only cytostatic effects also with doublet combinations (Fig. 2A, see days 6-13 and 21-24). However, fully supportive of our hypothesis, the triplet therapy treated cells were not able to escape the therapy during the follow up, and showed clear signs of cytotoxic response after initiation of the second dosing period (Fig. 2A, B). These results were validated across the heterogenous GB and GSC lines by using colony growth assays. Notably, regardless of importance of AKT-PDK axis in GB tumor growth,^42^ all cell lines, except for T98G, were resistant to combined AKT and PDK1-4 inhibition (DCA+MK-2206) (Fig. 2C, D). On the other hand, although NZ-8-061 was found to potentiate effects of MK-2206 or DCA to some degree across the cell lines, the triplet therapy (NZ-8-061+DCA+MK-2206) was again the only drug combination that was found effectively eradicating all GB and GSC lines, and without notable effects on fibroblasts (Fig. 2C, D). Fully validating PP2A reactivation as the mechanism inducing the synergistic drug interaction also in the context of triplet therapy, PME-1 inhibition induced synergism with combination of MK-2206 and DCA (Fig. S2C).

**Figure 2.**
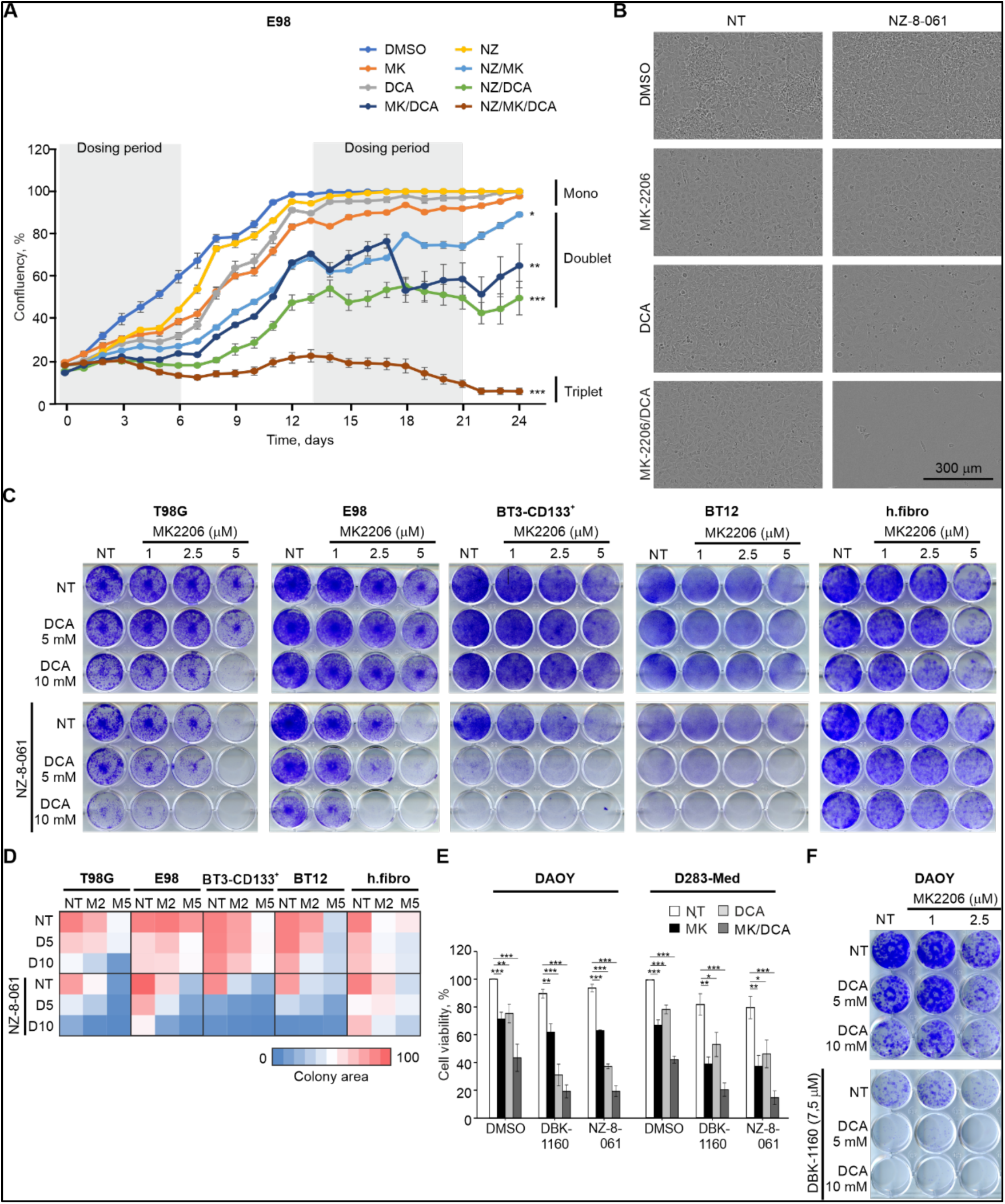
Triplet combination of NZ-8-061 with DCA and MK-2206 exerts a synergistic cytotoxic effect in molecularly heterogeneous GB and MB cell lines. **A)** Proliferation of E98 cells treated with DMSO, 7 µM MK-2206 (MK), 20 mM DCA, 10 µM NZ-8-061 (NZ) alone or in doublet or triplet combinations. Data as mean ±SEM (n = 6–12 wells per condition). *P<0.05, **P<0.01, ***P<0.001, Kruskal-Wallis test. **B)** Representative pictures of E98 cells from (A) at day 24. **C)** Representative images of colony growth assay in T98G, E98, BT3-CD133^+^, BT12 and fibroblasts under triplet combination treatment as indicated. **D)** Heat map representation of quantified colony growth assay data in the indicated cell lines treated with MK-2206 (MK; 2.5 and 5 µM), DCA (D; 5 and 10 mM) or NZ-8-061 alone or in doublet or triplet combination. Human fibroblasts were used as a negative control cell line. n=2 independent experiments. **E)** Cell viability in DAOY and D283-Med cells treated with DMSO, 8 µM DBK-1160 or 10 µM NZ-8-061 alone or in combination with 5 µM MK-2206 (MK), 20 mM DCA, or MK+DCA for 72 h. *P<0.05, **P<0.01, ***P<0.001, Student’s *t*-test. **F)** Representative images of colony growth assay in DAOY cells under the triplet combination as indicated.

Medulloblastoma (MB) is another brain tumor in which AKT and PDK kinase inhibitors have been proven clinically ineffective.^43^ Therefore, we studied whether the results above could be expanded from GB to MB. Reassuringly, when tested on two MB cell models, DAOY and D283-Med, representing SHH subtype and Group 3, respectively, we observed similar synergistic drug interaction between MK-2206, DCA and SMAPs (NZ-8-061 and DBK-1160) as across the GB cell lines (Fig. 2E). In addition, in colony growth assay in DAOY cells, we confirmed that combination of AKT and PDK inhibition was not sufficient for potent cytotoxicity, whereas combination with SMAP DBK-1160 resulted in very potent SL phenotype (Fig. 2F).

Collectively, these results demonstrate the brain tumor cells can escape kinase/PP2A targeting doublet combinations but cannot escape the triplet targeting of AKT, PDK1-4 and PP2A.

### The triplet therapy blunts therapy-induced signaling rewiring and potentiates apoptosis induction

Fully consistent with the therapy-induced signaling rewiring hypothesis behind inefficacy of kinase inhibitor therapies,^1,8,10^ we found that while MK-2206 efficiently inhibited the AKT S473 phosphorylation, it simultaneously enhanced phosphorylation of a direct mitochondrial PDK1-4 target PDHE1α (Pyruvate Dehydrogenase E1 Subunit Alpha 1)^40^ (Fig. 3A-C). In contrast, inhibition of PDK by DCA completely abolished phosphorylation of PDHE1α S300, but enhanced phosphorylation of AKT in T98G cells (Fig. 3A-C). However, combination of MK-2206 and DCA was able to shut-down phosphorylation of both proteins across all cell lines (Fig. 3A-C). NZ-8-061 treatment instead affected AKT and PDK signaling in very heterogeneous manner, depending on the kinase inhibitor combination, and the cell line. In other cell lines except for T98G, DCA+NZ-8-061 combination inhibited AKT S473 phosphorylation, but instead resulted in less efficient PDHE1α S300 inhibition than with DCA alone (Fig. 3C). On the other hand, NZ-8-061 did rescue the compensatory PDHE1α S300 phosphorylation induced by MK-2206. NZ-8-061 also expectedly inhibited AKT phosphorylation across the cell lines, but very interestingly also synergized with DCA in AKT inhibition (Fig. 3A-C).

**Figure 3.**
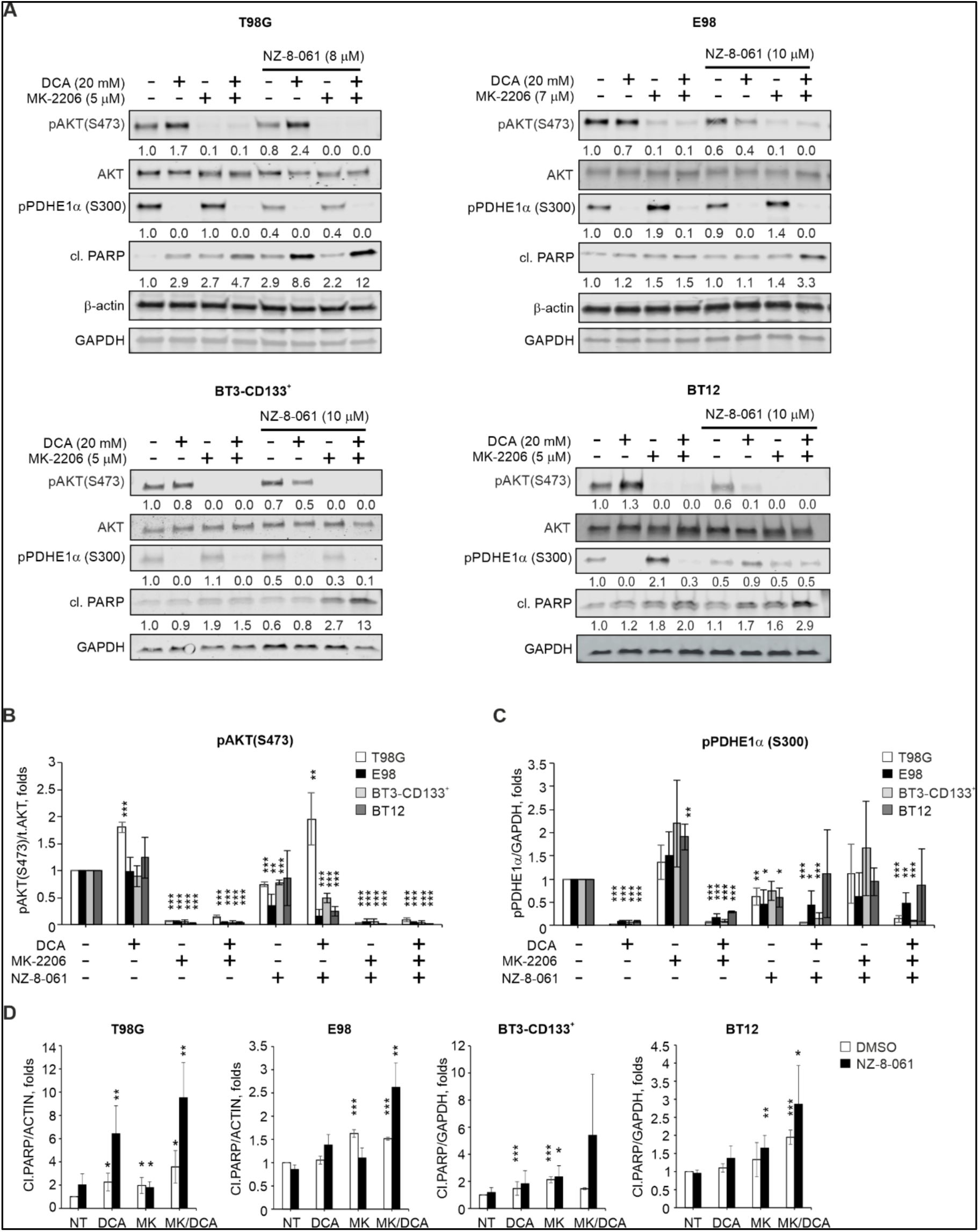
Inhibition of drug-induced signaling rewiring and apoptosis sensitization by the triplet therapy. **A)** Immunoblot assessment of phosphorylated AKT (S473), phosphorylated PDHE1α (S300), and cleaved PARP after treatment with DCA, MK-2206 or NZ-8-061 alone or in doublet or triplet combination for 24 h in T98G, E98, BT3-CD133^+^ and BT12 cells. Normalized quantifications from (A) for **B)** phosphorylated AKT (S473), **C)** phosphorylated PDHE1α, and **D)** cleaved PARP. *P<0.05, **P<0.01, ***P<0.001, Student’s *t*-test.

To correlate these findings to the apoptotic potential of the combination therapies, we examined PARP cleavage from the same cellular lysates. The data reveals that neither total shutdown of AKT and PDK axis (MK-2206+DCA) or NZ-8-061 at doses that synergize in drug combinations (Fig. 3B, C), was sufficient for maximal apoptosis induction in any of the studied GB cell lines (Fig. 3D). However, the highest apoptotic response was consistently seen across all cell lines upon the triplet therapy treatment (Fig. 3D). DAOY MB cells also displayed similar therapy-induced signaling rewiring between AKT and PDK pathways, but combination with DBK-1160 blunted the rewiring and resulted in potent apoptosis induction (Fig. S2D).

These observations confirm prevalent therapy-induced signaling rewiring and heterogeneity in the combinatorial drug responses across the GB cells. Importantly, the triplet therapy was found to inhibit therapy-induced signaling rewiring, and thereby convert cytostatic kinase inhibitor responses to cytotoxic effects across GB cells.

### Triplet therapy inhibits mitochondrial OXPHOS and primes to BH3 protein-mediated apoptosis

The results above revealed that pharmacological PP2A reactivation can impact mitochondrial PDK signaling. To assess basal cellular metabolic activity, T98G cells were exposed to either MK-2206, DCA or NZ-8-061 alone or in combination, and Seahorse XF Cell Mito Stress Test was applied. As expected, DCA alone increased ATP production, as it reactivates the OXPHOS in the mitochondria (Fig. 4A).^40,44^ On the contrary, MK-2206 reduced ATP production and mitochondrial-linked respiration (Fig. 4A). Interestingly, NZ-8-061 used at SL inducing non-toxic concentration had a broad-spectrum effect on mitochondrial metabolism. Especially interesting drug interaction was inhibition of DCA-induced OXPHOS (Basal, Maximal, and Spare), indicating that PP2A reactivation can prevent compensatory mitochondrial survival mechanism.

**Fig. 4.**
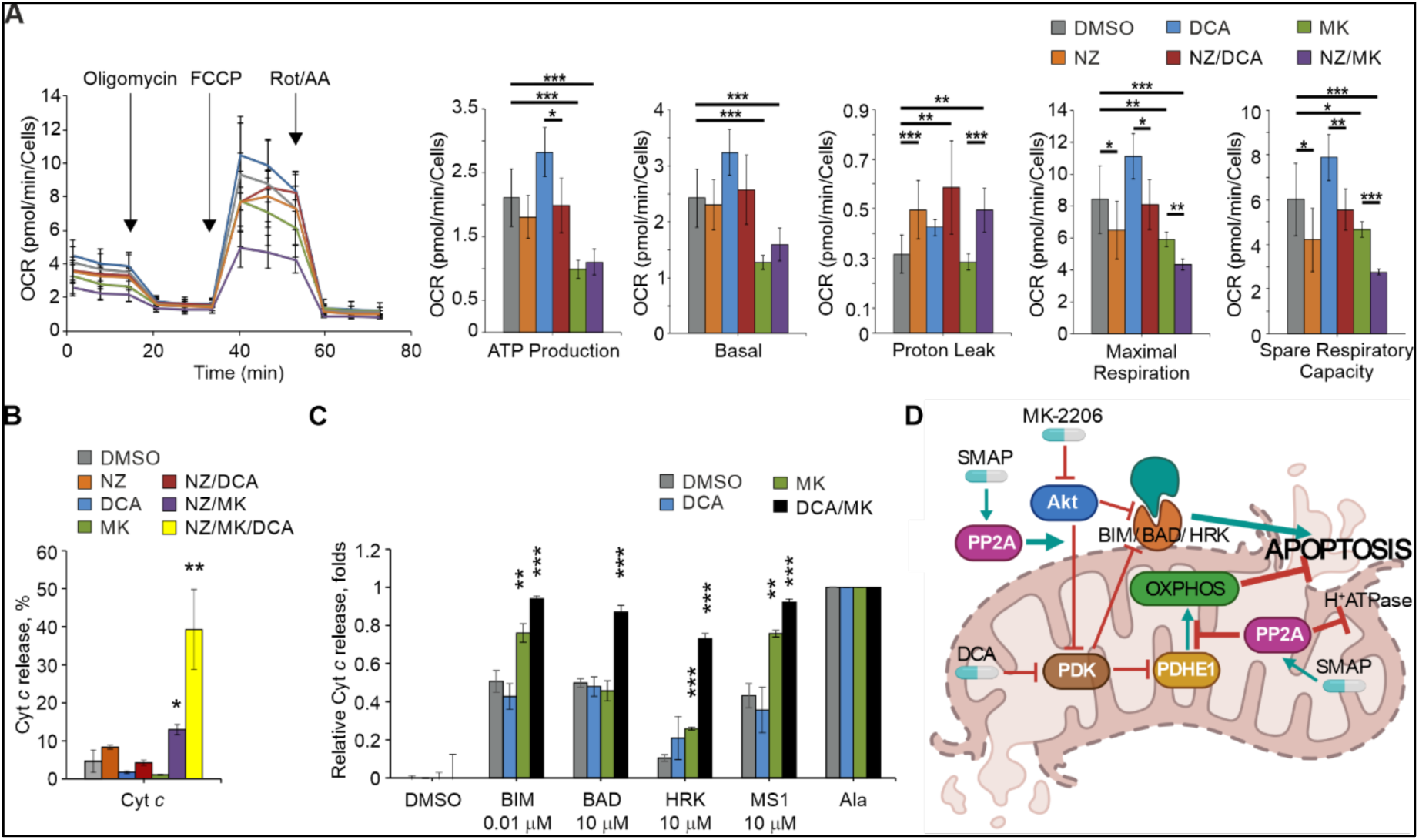
Triplet therapy inhibits mitochondrial OXPHOS and primes to BH3 protein-mediated apoptosis. **A)** Mitochondrial stress test Seahorse profile and mitochondrial parameters in T98G cells treated with 10 mM DCA or 7 µM MK-2206 (MK) alone or in combination with 10 µM NZ-8-061 (NZ) for 24 h. *P<0.05, **P<0.01, ***P<0.001, Student’s *t*-test. **B)** Cytochrome *c* release from T98G cells treated with 5 µM MK-2206 (MK), 20 mM DCA or 8 µM NZ-8-061 (NZ) alone or in doublet or triplet combination for 24 h. *P<0.05, **P<0.01, ***P<0.001, Student’s *t*-test. **C)** Priming of T98G cells to apoptosis induction by indicated BH3 peptides. T98G cells treated with 5 µM MK-2206 (MK), 20 mM DCA alone or combination for 24 h. *P<0.05, **P<0.01, ***P<0.001, Student’s *t*-test. **D)** Schematic illustration of mitochondrial mechanisms for the triplet therapy-induced apoptosis. Inhibition of PDK1-4 induces compensatory OXPHOS but this is blunted by SMAP treatment which additionally induces mitochondrial membrane proton leakage. PDK1-4 and AKT inhibition synergizes on BH3-meidated apoptosis priming and SMAP treatment inhibits signaling rewiring between the kinases. Whereas in response to doublet drug combinations cells can induce some compensatory survival mechanism, these are simultaneously inhibited in triplet therapy treated cells resulting in terminal apoptosis induction.

NZ-8-061 alone, and in combination with MK-2206, also profoundly increased proton leak indicating for mitochondrial membrane damage (Fig. 4A). Further, in line with only cytostatic effects with mono-and doublet therapies there was no mitochondrial cytochrome *c* release by any single drug treatments, or with doublet combinations (Fig. 4B). In contrast, the triplet therapy induced strong cytochrome *c* release (Fig. 4B). As cytochrome *c* release is controlled by BH3-only proteins on the outer mitochondrial membrane,^45^ we clarified the functional interaction between cytoplasmic AKT, and mitochondrial PDK1-4 kinases on regulation of mitochondrial apoptosis by dynamic BH3 profiling.^26,27^ BH3 profiling revealed a limited impact on apoptotic priming by PDK1-4 inhibition, but a marked increase in the cells’ susceptibility towards BIM, HRK, and MS1 mediated cytochrome *c* release when AKT was inhibited (Fig. 4C). Notably, there was a marked enhancement and broadening of BH3-mediated apoptosis priming when AKT and PDK1-4 were co-inhibited, providing an additional explanation for their synergistic pro-apoptotic effect (Fig. 4D). Results related to the impact of triplet therapy on BH3 profiling were inconclusive presumably due to high apoptotic activity (data not shown).

Collectively, these data reveal the mechanistic basis for the high apoptotic activity of the triplet therapy. We conclude that the cytotoxicity is mediated by PP2A-elicited inhibition of the compensatory OXPHOS and induction of inner mitochondrial membrane proton leakage, combined with synergism between MK-2206 and DCA on BH3 priming.

### Validation of therapeutic potential of the triplet therapy in orthotopic GB and MB models

*In vivo* relevance of the results was investigated in subcutaneous and intracranial models using both E98 GB cells and DAOY MB cells. First, we wanted to provide *in vivo* validation to AToMI screening results^24^ that the SL effects of SMAPs with UCN-01 can be recapitulated by combination of AKT and PDK inhibition. As UCN-01 does not cross the BBB, these first *in vivo* experiments were performed using subcutaneous xenografts, and instead of NZ-8-061, we used DBK-1160 as a SMAP due to its better pharmacokinetic profile based on our previous studies (data not shown). Fully validating the *in vitro* results, the orally dosed triplet therapy (DBK-1160+MK-2206+DCA) was equally efficient, or even superior to combination of DBK-1160 and UCN-01 (Fig. 5A, B). The robust *in vivo* antitumor effect of the triplet therapy in DAOY model was readily seen also when comparing the sizes of the excised tumors (Fig. 5C).

**Fig. 5.**
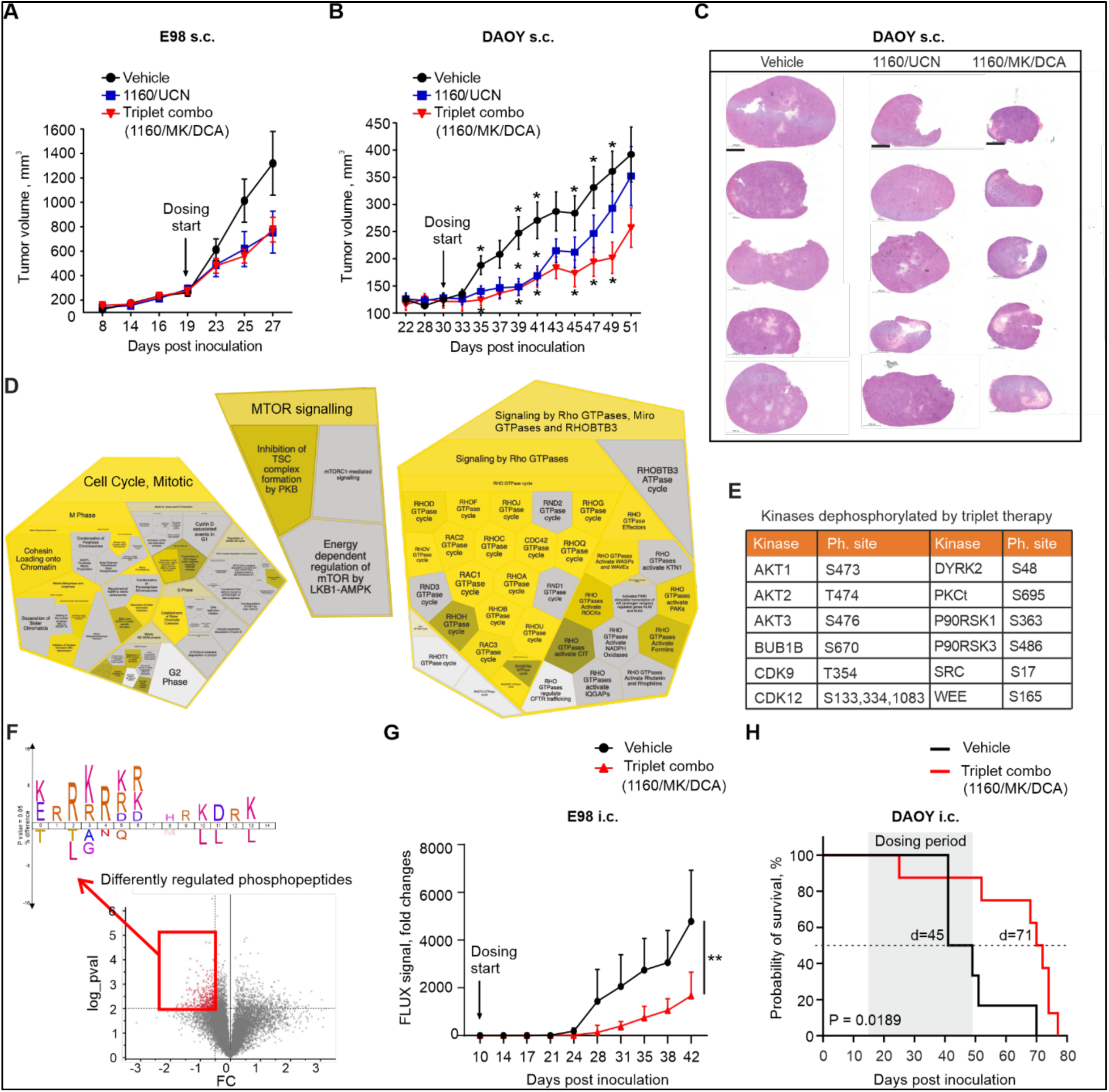
Validation triplet combination therapy in vivo. Quantification of tumor volume from E98 **(A)** and DAOY **(B)** s.c. tumors in mice treated with DBK-1160 (1160; 100 mg/kg twice a day) and UCN-01 (UCN; 3 mg/kg once a day) or MK-2206 (MK; 100 mg/kg every second day) and DCA (100 mg/kg twice a day), or vehicle control. Each group had n=8 mice in E98, n=10 mice in DAOY experiments. *P<0.05, two-way ANOVA test. **C)** Representative images of H&E staining DAOY s.c. tumors from (B, n=5). Scale bar, 1000 µm. **D)** Reactome processes based on significantly (*P<0.05) regulated phosphopeptides from triplet therapy treated DAOY xenografts in (B). **E)** Kinases dephosphorylated by the triplet therapy in DAOY xenografts from (B). **F)** Volcano plot showing differentially regulated phosphopeptides from (B). Icelogo kinase motif enrichment analysis from the dephosphorylated peptides (in red) (**P≤0.01, log2FC≤-0.5) revealed enrichment of canonical AKT sites (R-x-R-x-x-S/T and R-x-x-S/T). **G)** Bioluminescence follow up of an orthotopic E98 glioblastoma tumor comparing the vehicle or triplet combination therapies (DBK-1160 (100 mg/kg twice a day) + MK-2206 (100 mg/kg every second day) + DCA (100 mg/kg twice a day)). Mean ±SEM. n=10 mice per group. **P<0.01, Student’s *t*-test. **H)** Kaplan–Meier survival analysis of xenograft orthotopic DAOY model treated with triplet combination (DBK-1160 (100 mg/kg twice a day) + MK-2206 (100 mg/kg every second day) + DCA (100 mg/kg twice a day)). Vehicle n=6, Triplet combo n=8 mice. *P<0.05, Mantel-Cox test.

To molecularly profile the triplet therapy effect in the treated tumors, the vehicle and the triplet therapy treated tumors (n=5 for both) were subjected to MS-phosphoproteomics analysis. Upon filtering the data for those phosphopeptides there were quantifiable from at least three tumors per group, and with FDR of 5% for significance of the difference in phosphopeptide expression between the groups (Table S1), the Reactome pathway analysis validated the impact triplet therapy on both apoptosis and cell cycle, but on the other hand revealed a very strong enrichment of targets involved in “Signaling by Rho GTPases” (Fig. 5D, S3, Table S2). Furthermore, fully consistent with our model that efficient therapy response in brain tumors requires wide-spread kinase inhibition, we found inhibition of phosphorylation of several kinases from the triplet therapy treated tumors (Fig. 5E, Table S1). Notably, among those were inhibition of the phosphorylation of the activation loop of AKT1, 2 and 3 (Fig. 5E), which together with enrichment of mTOR signaling based on phosphopeptide data (Fig. 5E, Table S1), perfectly supports our mechanistic data demonstrating importance of the shutdown of rewiring to AKT signaling (Fig. 3). Inhibition of AKT signaling was evident also based on kinase target motif enrichment analysis where canonical AKT target motifs (R-x-R-x-x-S/T and R-x-x-S/T) were clearly enriched in the phosphopeptides downregulated by the triplet therapy (Fig. 5F). Beside AKT, among the dephosphorylated kinases were also transcriptional elongation promoting kinase CDK9, that is essential for brain tumor-initiating cells,^46^ and a synergistic drug target with SMAPs.^21^ Interestingly, but consistent with the therapy-induced non-genetic signaling rewiring, we also identified a number of phosphopeptides upregulated in triplet therapy treated tumors (Fig. 5F upper right corner, Table S1). Related to kinase signaling, we noticed that several kinases involved in the pro-apoptotic JNK and p38 MAPK signaling were hyperphosphorylated in the treated tumors (Fig. S4), and both JNK1 and JNK2 were among the top enriched kinase target motifs based on NetworKIN analysis (Fig. S4A). As both JNK and p38 are involved apoptosis regulation by BH3 proteins,^47^ these data provide a plausible link between the proposed mechanism for triplet therapy induced brain tumor cell killing, and the observed in vivo therapeutic effects.

Finally, the triplet therapy was tested in intracranial model with luciferase-expressing E98 cells that carry characteristics of GSCs and have very infiltrative growth pattern *in vivo*.^22^ In addition to these faithful human GB characteristics, E98 cells displayed indistinguishable triplet therapy response as compared to patient derived GSC cell lines *in vitro* (Fig. 2). Importantly, we observed significant inhibition of tumor growth by orally dosed triplet therapy initiated upon appearance of detectable tumors at day 10 (Fig. 5G). For DAOY cells, we relied on mouse survival as the end-point measurement of the therapy effect, since no tumor growth visualization approaches were available for these tumors. Remarkably, more than 50% of the vehicle treated mice died during the therapy, whereas in the triplet therapy group only one mouse had to be sacrificed due to neurological symptoms (Fig. 5H). Following cessation of therapy after 30 days, due to local regulations, we observed a significant increase in mouse survival in the triplet therapy group, associated with 26-day prolongation of the median probability of survival (Fig 5H). No obvious toxicities were observed during triplet therapy treatment periods in either subcutaneous or intracranial models (Fig. S5). However, as expected, the SMAP treatment resulted in reversible increase in liver weight, as has been reported earlier.^30^

## DISCUSSION

Kinase inhibitor resistance of brain tumors is a notable unmet clinical challenge.^4,17,43^ Considering that hyperactivated kinase signaling is one of the hallmarks of GB,^3,11^ clinical resistance of GB to kinase inhibitors constitutes also a clear mechanistic enigma. There is a strong theoretical basis for synergistic activities of simultaneous kinase inhibition and phosphatase activation in phosphorylation-dependent cancers,^10,20^ but the therapeutic impact of such combinatorial approach in brain cancers has been thus far unclear. Here we demonstrate that heterogeneous GB and MB cell lines have astonishing capacity to escape combination of inhibition of one kinase and PP2A reactivation. However, our results clearly demonstrate that kinase inhibitor tolerance in brain cancers can be overcome by targeting simultaneously three phosphorylation-dependent signaling nodes: AKT, PDK1-4 and PP2A.

MKIs provide an attractive approach to simultaneously inhibit several oncogenic kinases, and some MKIs (e.g., Sunitinib, PKC412), are clinically used as cancer therapies.^48^ However, similar to more selective kinase inhibitors, all tested MKIs have thus far failed in GB clinical trials.^4^ To better understand GB relevant STS target kinases, we developed the AToMI approach^24^ and found several kinases which synergized with PP2A reactivation by either PME-1 inhibition or by SMAPs. Notably, the kinases which synergized with PP2A reactivation represent the commonly hyper activated pathways in GB. For example, AKT pathway is one of the most dysregulated pathways in GB whereas PDK1-4 kinases have a critical role in GB mitochondrial glycolysis. However, AKT and PDK1-4 targeting monotherapies have failed to demonstrate significant survival effects in clinical trials for GB.^40,49-51^ Our subsequent kinase inhibitor combination experiments, using inhibitors at doses that inhibit their target kinases, validate the ineffectiveness of even combinatorial targeting of AKT and PDK1-4 in eradicating heterogeneous GB cell lines. This was regardless that the therapy-induced signaling rewiring between AKT and PDK pathways was abolished when MK-2206 and DCA were combined. Regardless of importance of AKT-PDK axis in GB tumor growth,^42^ these results challenge the concept that targeting of AKT-PDK1-4 axis would be sufficient for GB therapy. Instead, our data undeniably demonstrate the triplet therapy including, in addition to AKTi and PDK1-4i, also PP2A reactivation eradicates all tested cell models.

Collectively, our data identity a strategy for killing of heterogeneous brain tumor cells based on triplet kinase/phosphatase targeting of critical GB and MB signaling nodes. Notably, our results are relevant across heterogeneous GB and MB models including patient-derived GSCs. Based on our results, the uniform kinase inhibitor resistance observed in GB and MB clinical trials,^4^ could be to significant extent contributed to non-genetic PP2A inhibition by PME-1.^32^ In this regard, the current results encourage future brain tumor clinical trials in a significant proportion of brain cancer patients with low tumor expression of PME-1^23^ with combinations of clinically tested AKT and PDK1-4 inhibitors.^16,49,51^ Importantly, diagnostic definition of PME-1 status would greatly simplify biomarker-based analysis of PP2A activity in brain tumors because it sidesteps the need for analyzing all the possibly relevant PP2A subunits. Finally, our results strongly indicate that rapidly developing PP2A reactivation therapies^20^ will constitute an attractive future therapy option for brain tumors when combined with multi-kinase inhibition.

## Funding

Sigrid Juselius Foundation (J.W.); K. Albin Johanssons Foundation (J.W.); Aamu Pediatric Cancer Foundation (201900013, O.V.D.); Turku Doctoral Programme of Molecular Medicine (J.M.); Academy of Finland (T.A.); Finnish Cancer Foundation (180157, J.W.).

### Acknowledgements

Authors want to acknowledge Biocenter Finland infrastructures, especially Turku Proteomics Facility, and Genome Editing Core at Turku Bioscience Centre, High Throughput Biomedicine Unit at Institute for Molecular Medicine Finland, Turku Center for Disease Modelling at University of Turku. Personal acknowledges to Nikhil Gupta for help with nucleofection technique, William Eccleshall and Mung Kwan Long for help with Seahorse experiments, Kari Kurppa for expert help with Incucyte experiments and for helpful discussions, Johanna Ivaska for useful comments on the manuscript, Taina Kalevo-Mattila for excellent technical support as well as the entire Turku Bioscience personnel for excellent working environment.

## Authorship

Conception and design: O.V.D., J.M., A.K., J.W. Development of methodology: O.V.D., R.H., O.K., J.W. Experimental work: O.V.D., J.M., X.Q., M.T., C.S-F., K.W., R.K., M.P. Bioinformatic analysis: M.J., L.Y., O.K., T.A. In vivo work: O.V.D., J.M., R.H. Resources: M.O. Writing: O.V.D., J.M., T.A., J.W.

## Data and materials availability

All data associated with this study are present in the paper or the Supplementary Materials.

